# An Interpretable Framework to Characterize Compound Treatments on Filamentous Fungi using Cell Painting and Deep Metric Learning

**DOI:** 10.1101/2023.08.24.554566

**Authors:** Laurent Lejeune, Morgane Roussin, Bruno Leggio, Aurelia Vernay

## Abstract

The cell painting microscopy imaging protocol has recently gained traction in the biology community as it allows, through the addition of fluorescent dyes, to acquire images that highlight intra-cellular components that are not visible through traditional whole-cell microscopy. While previous works have successfully applied cell painting to mammalian cells, we devise a staining protocol applicable to a filamentous fungus model. Following a principled visual inspection and annotation protocol of phenotypes by domain-experts, we devise an efficient, robust, and conceptually simple image analysis strategy based on the Deep Cosine Metric Learning paradigm that allows to estimate phenotypical similarities across different imaging modalities. We experimentally demonstrate the benefits of our pipeline in the tasks of estimating dose-response curves over a wide range of subtle phenotypical variations. Last, we showcase how our learned metrics can group image samples according to different modes of action and biological targets in an interpretable manner.

## 1. Introduction

Pathogenic fungi are known to be the dominant cause of plant diseases and incur paramount economical damages to commodity crops [10]. Hence, crop protection strategies aim to discover relevant fungicides to prevent these, by leveraging findings pertaining to various scientific domains such as biology, chemistry, and genetics, to name a few [8].

In this context, a past contribution combined microscopy imaging with computer vision and machine learning techniques, and showed that compound treatments can induce distinctive phenotypical changes to a filamentous fungi, which in turn are crucial information that allow to raise a well-informed hypothesis on the possible biological targets of a compound [18].

In parallel, an emerging multiplex microscopy imaging technique called cell painting has recently gained traction[3]. At its core, cell painting consists in adding, as part of a plate assay, fluorescent dyes known to bind to specific extra and intra-cellular compartments, while appropriate filters are setup in a microscope to separate the fluorescent emissions at different wavelengths. As a result, one obtains for each filter an additional image-channel. Setting up a painting protocol requires the careful optimization of many parameters: to name a few, the selection of labelling reagents able to penetrate the fungal wall, the type of dyes, their combinations and concentrations, and the selection of agents used to wash plates [6].

In the present work, we apply the cell painting protocol to the filamentous fungus model *Botrytis cinerea* [1], a widespread plant pathogen, through an adequate staining protocol, which, to the best of our knowledge, has not been done in the past. We further devise an appropriate computer vision and machine learning strategy to investigate specific intra-cellular morphological impacts of compound treatments. In particular, we contribute a simple yet effective solution based on Deep Cosine Metric Learning that allows to (1) infer measures of biological activity of compound treatments by means of dose-response curves, as well as (2) gain insights into the biological targets of a treatment through cluster analysis. Importantly, we put an emphasis on producing results that are easy to interpret by domain-experts. In particular, we learn a set of metrics-spaces that correspond to biologically meaningful categories, each focusing on one intra-cellular component (lipid droplets, cell wall, …) or whole-cell morphology (type of germination).

The present work is organized as follows. In the related works section, we give an overview of past contributions related to computer vision and machine learning methodologies that target compound treatment characterization using cell painting image data. In the methods section, we briefly describe our imaging protocol, and develop our machine learning solution. We then lead in-silico experiments where we compare the proposed method with baseline methods for dose-response curve estimation and cluster analysis.

## 2. Related Works

Imaging has been used extensively across many scientific disciplines, and in particular in biology, as a mean to identify relevant phenotypes of organisms [13]. A popular approach consists in extracting a morphological profile on each imaged cell, i.e. a set of pre-defined visual features assumed to be informative for downstream tasks[29].

A popular choice in this context is Cell-Profiler, a feature extraction framework that allows to extract visual features specifically targeted to cellular microscopy images of mammalian cells [3] Previous works have leveraged this framework in the frame of cell painting plate assays to extract 1500 features per imaged cell [21, 17], and effectively use these to identify genetic perturbations and predict modes of action (MoA)[14]. Despite being a potent approach in the latter context, this pipeline assumes that the object of interest, i.e. cells, form well defined blobs. This assumption is largely inadequate in the context of filamentous fungi, where the object of interest is much more complex and importantly, not compact.

Fortunately, more robust and flexible alternatives exist, namely those relying on Deep Convolutional Neural Networks (CNN)[5]. In particular, models relying on CNNs typically take as input the entire image and produce relevant outputs (features, predictions, …) through the optimization of an objective function. Importantly, these models implicitly learn to detect objects of interest within an image, and aggregate relevant features at different scales [28]. In the context of cell painting imaging, [4] leverages a CNN to extract features using morphological and phenotypical similarities of lung tissues that correspond to the same treatment, while as a downstream task, authors group variants of the same gene while discarding its mutations. In [14], authors provide a comparative benchmark of feature extraction methods applied to the prediction of chemical perturbations from treated mammalian cells. In particular, their experiments show that methods relying on Cell-Profiler features can be surpassed on these tasks through deep network models, while bypassing the cumbersome segmentation step.

## 3. Methods

We start this section by giving a brief description of the cell painting protocol used throughout our experiments. Next, we describe our exploration and annotation procedure, by which a domain-expert selects among a wide range of images a set of representative samples that show biologically relevant phenotypes. We then describe our strategy based on deep cosine metric learning, where we train a model that learns a set of metric spaces, where each space effectively clusters a given biologically relevant category of phenotypes, while each cluster represents a distinct phenotype. Last, we emphasize how the cosine softmax training paradigm, by design, provides a distance measure applicable to dose-response estimation. We show in Fig. 1 an overview of our pipeline.

**Figure 1.**
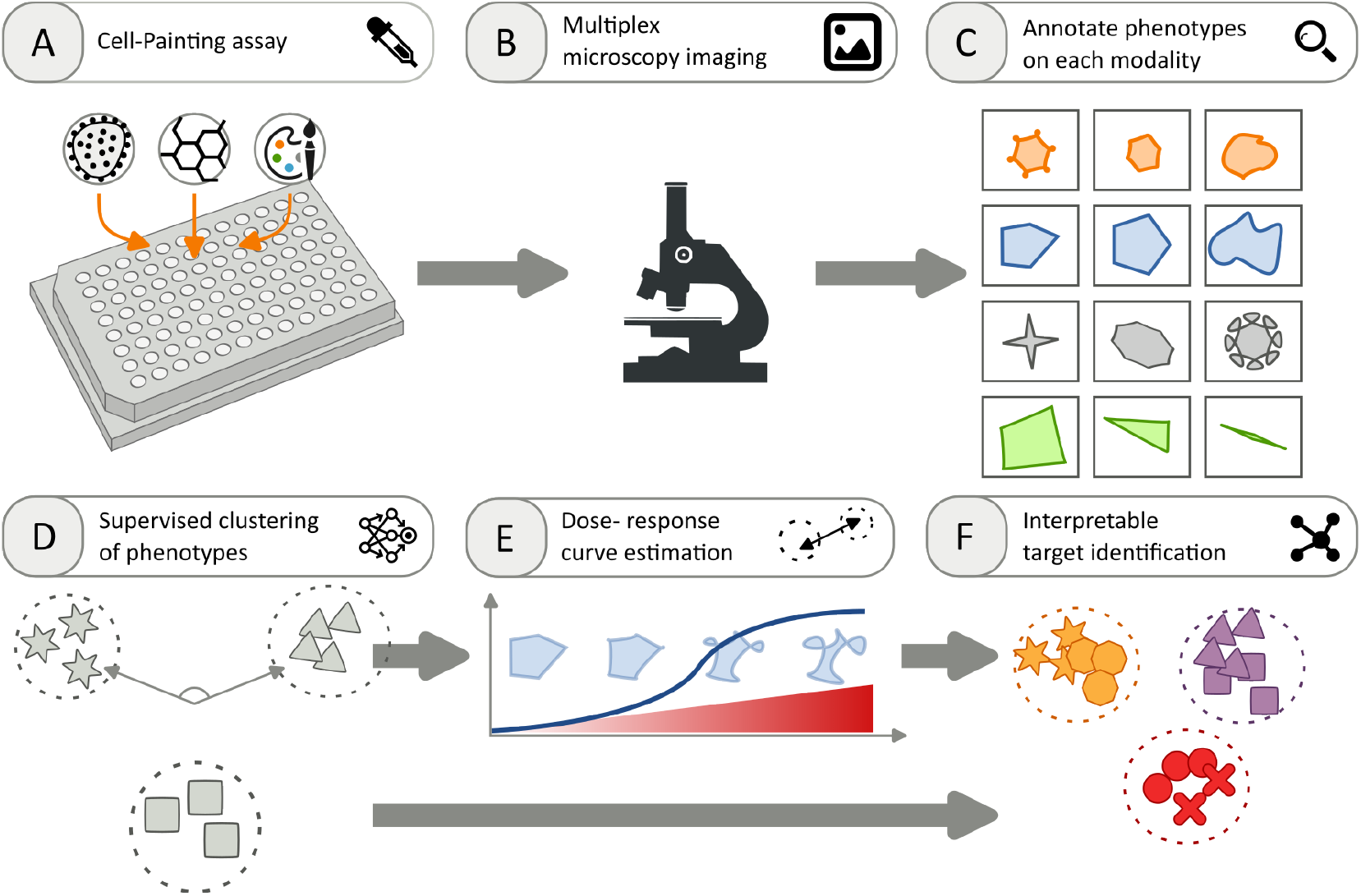
Overview of our contributions. (**A**) We implement a cell painting imaging protocol adequate to a filamentous fungus so as to capture microscopy images of different intra-cellular components not observable with traditional microscopy. (**B**) We acquire, for different compounds and doses, images on 3 fluorescent image channels, each emphasizing a specific component, i.e. lipids, cell wall, and apical cell wall. (**C**) A domain-expert explores images and provides qualitative labels on each modality, where labels correspond to phenotypes of an intra-cellular component, or a global morphological configuration. All phenotypes are organized in biologically meaningful categories (**D**) We optimize a supervised metric learning model that clusters phenotypes in each category independently. (**E**) We test different compounds at increasing concentrations and estimate, for each category, a dose-response curve. (**F**) We identify biologically active samples from the previous phase, and leverage the same model to perform a cluster analysis according to biological targets.

### 3.1. Imaging protocol, exploration, and annotation procedure

We implement a cell painting protocol for the filamentous fungus *Botrytis cinerea*, and select 3 fluorescent dyes: Nile Red, Calcofluor (CFW), and Wheat Germ Agglutinin (WGA) to stain lipids [23], cell wall, and apical cell wall [16], respectively. We give further technical details in Tab. 1. As microscope, we use a MicroXL (Molecular Devices) with magnification 40x, which provides on each imaged location a field of view of 350 *×* 350*μm*.

**Table 1.** Set of dyes used in our cell painting protocol.

| Dye | Reference | Excitation [nm] | Emission [nm] | Exposition [ms] | Concentration [$\mu\text{g/ml}$] |
| --- | --- | --- | --- | --- | --- |
| Nile Red | Sigma 19123 | 530 | 635 | 30 | 5 |
| CFW | Sigma 3543 | 380 | 475 | 50 | 6 |
| WGA | Invitrogen W11261 | 488 | 549 | 50 | 0.5 |

In order to explore and capture a broad variety of phenotypes on our filamentous fungus model, we expose it to 24 different compound treatments as well as negative control conditions (DMSO) on 5 plate assays, each containing 96 wells, so as to obtain *∼* 13800 images. The compounds and their concentrations were selected by domain experts so as to cover representative MoAs and targets described by the Fungicide Resistance Action Committee (FRAC) [9]. After visual inspection, we retain *∼* 8380 images across all modalities. Next, we assign a label to each phenotype and group them into categories to obtain 7, 6, 4, and 7 phenotypes for category lipid, cell wall, apical cell wall, and morphology, respectively^1^.

We show in Fig. 2 a subset of images from our training dataset along with a code-name, and give textual descriptions of corresponding phenotypes in Tab. 2.

**Figure 2.**
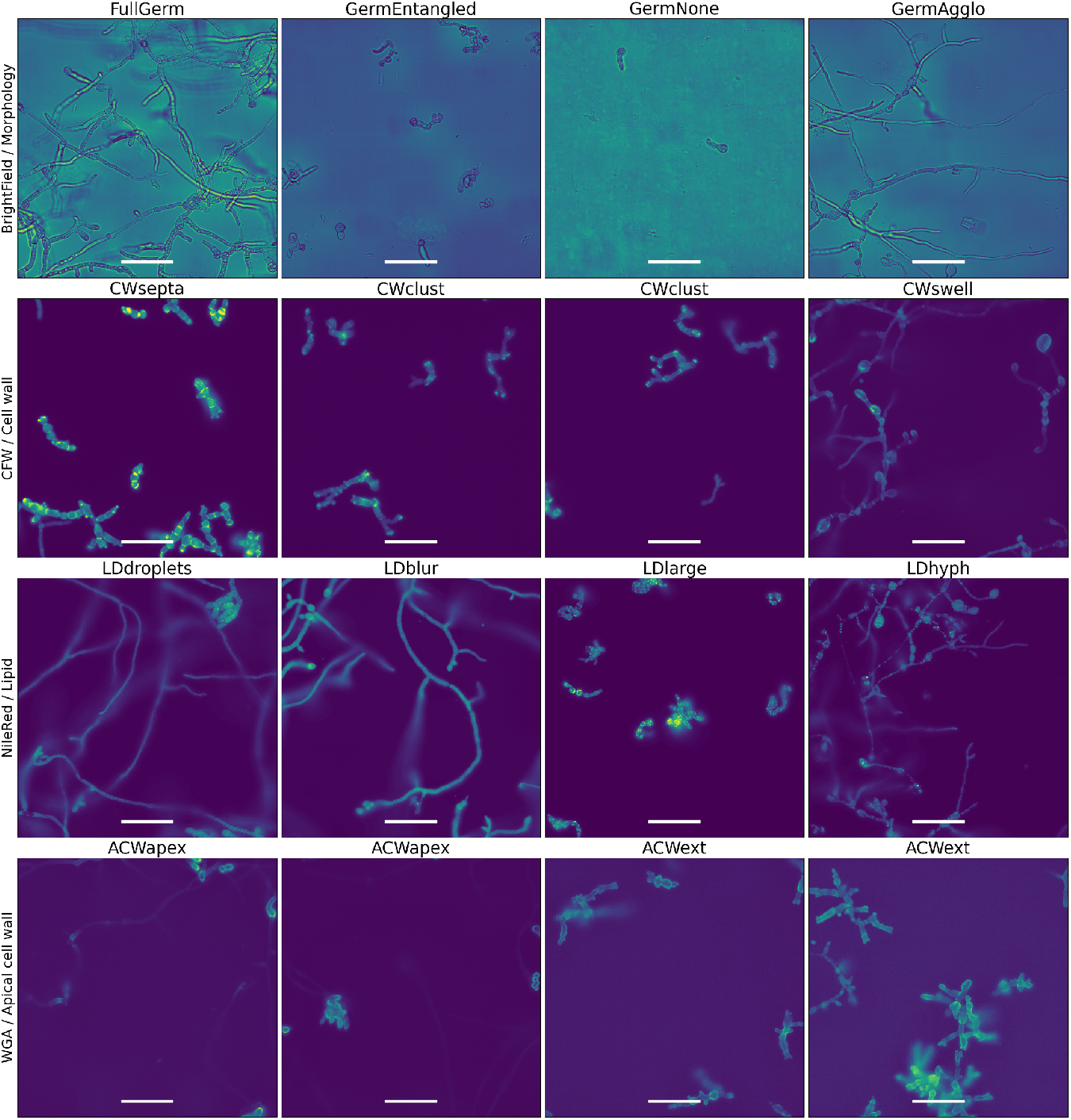
Example cell painting images. Each row correspond to a phenotypical category, while columns correspond to phenotypes. Scalebar: 70*μm*.

**Table 2.** Description of a subset of annotated phenotypes for each category.

| Category | Dye/Modality | Phenotype |  |
| --- | --- | --- | --- |
|  |  | Code | Description |
| Lipid | Nile-Red | LDdroplets | Sharp and granular lipid droplets |
|  |  | LDnone | Homogeneous staining within spores and hyphae |
|  |  | LDlarge | Large lipid droplets |
|  |  | LDhyph | Homogeneous staining within spores, large blobs in hyphae |
| Cell wall | Calcofluor (CFW) | CWsepta | Sharp and granular staining at Septa and Apex |
|  |  | CWclust | Diffuse and intense signal |
|  |  | CWswell | Swelling patterns |
| Apical cell wall | Wheat Germ Agglutinin (WGA) | ACWapex | Sharp staining at apex |
|  |  | ACWext | Spatially extended stainings |
| Morphology | Transmitted Light | FullGerm | Full germination of mycelium |
|  |  | GermNone | No germination |
|  |  | GermEntangled | Entangled hyphae |
|  |  | GermAgglo | Agglomerate of spores |

### 3.2. Deep Cosine Metric Learning (DCML)

We now develop our proposed deep cosine metric learning method, which we name **DCML**, and emphasize its pecularities and relevance for the practical problems tackled in this work.

Let *C* = *{c*_*k*_|*k* = 0, …, *K −* 1*}* a set of *K* phenotypical categories, and 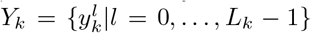 a set of *L*_*k*_ labels denoting distinct phenotypes that lie in category *c*_*k*_. Next, let 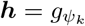*○ f*_*θ*_(*I*) *∈* ℝ ^*D*^ an embedding obtained by successively passing image *I* in a feature-extraction function *f*_*θ*_, followed by 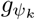, which further transforms the feature vector into a category-specific embedding that encodes distances. Last, we look to obtain for each phenotypical category *c*_*k*_ a *bounded* distance function *d*_*k*_(***h***_*i*_, ***h***_*j*_), that takes values 0 when ***h***_*i*_ and ***h***_*j*_ correspond to two images that show identical phenotypes, and 1 otherwise. The boundedness criteria, as will be shown in our experiments, is of practical interest when inferring dose-response curves.

Taking inspiration from [27], we solve the above metric-learning problem through the cosine softmax classifier approach. For conciseness, we give here a brief overview of the theoretical development, and refer interested readers to the original work.

Without loss of generality, we omit index *k* for the remaining of this section. In a standard softmax classification setting, one typically obtains probability scores for each class by applying a linear transformation parameterized by weights ***ω*** and offset ***b*** via

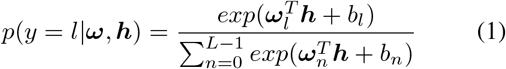

Interestingly, Eq. 1 corresponds to the posterior probability function of Gaussian densities *N* (***μ***_*l*_, **Σ**), where ***ω***_*l*_ = **Σ**^*−*1^***μ***_*l*_, and 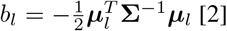.

Next, the parameterized encoder and linear layer are optimized by means of the cross-entropy loss:

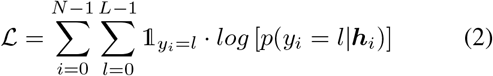

As noted in [27], a softmax classifier optimizes for Eq. 2 in a discriminative fashion, i.e. it aims at bringing the posterior *p*(*y* = *l*|***h***) as close as possible to the corresponding indicator function. Importantly, it does so by pushing embeddings ***h*** away from the decision boundaries *b*_*l*_, but not necessarily closer to their class centroids.

To circumvent this limitation, the cosine softmax classifier framework explicitly optimizes for class-centroids and embeddings in a joint manner. Concretely, the class-centroids and embeddings are *ℓ*_2_ -normalized, i.e. 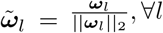, *∀l*, and 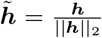. Finally, the cosine softmax classifier output probabilities are

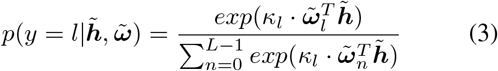

where *κ*_*l*_ is a scaling parameter. Similar to the standard softmax classification setting, we perform optimization using the cross-entropy loss (Eq. 2). Through a simple change of parameters and normalization, one can show through Baye’s rule that we explicitly enforce that the class-conditionals follow a von Mises distribution [19] centered on class centroids, i.e.

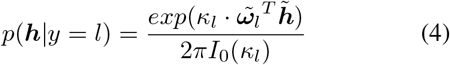

with *I*_0_ the modified Bessel function of the first kind of order 0. In words, the probability density in Eq. 4 peaks at 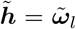, while the scaling parameter *κ*_*l*_ governs the rate of decay.

To summarize, through the cosine softmax classification framework, we optimize for embeddings that are appropriate for pairwise similarity measurements because embeddings form compact clusters around their class centroids, while class centroids are pushed away from each other so as to spread on the unit-sphere.

#### 3.2.1 Calibration, Dose-Response Curve Estimation, and Monotonicity Constraint

In the context of dose-response estimation, we wish to infer, given a test and a negative-control population, an interpretable measure of perturbation. By convention, a response-level of 0 corresponds to a compound that induces no perturbation, while a response-level of 1 corresponds to a condition where the cell incurs maximal perturbation. We now suggest a simple calibration step to convert cosine similarity scores to interpretable response-levels.

From our validation set, we estimate parameters *α*_*k*_, each corresponding to the expected cosine of dissimilar pairs for category *k*. While an intuitive approach would be to select these as the arithmetic mean of cosine similarities of dissimilar pairs, we observed that the empirical distributions often exhibit a skew, and rather chose the value that maximizes the latter (see Fig. 3).

**Figure 3.**
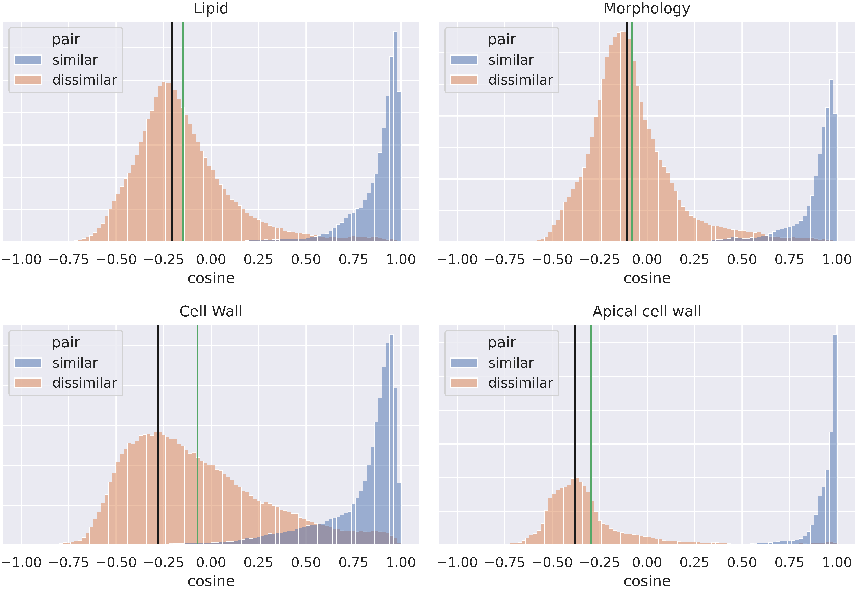
Cosine similarity distributions of similar and dissimilar image-pairs taken from a validation set on each category. The black and green vertical lines are the maximum peak and the mean, respectively.

Next, we define our bounded category-specific distance function as

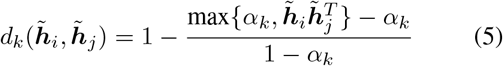

We let *ℌ* and *ℌ*_0_ two sets of embeddings that correspond to the test condition and the negative-control condition image-sets, respectively. Finally, we obtain the responselevel *r*_*k*_ according to category *k* by averaging distances on all pairs of embeddings in *ℌ × ℌ*_0_, i.e.

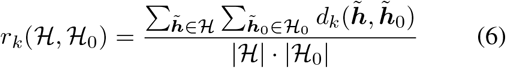

Following guidelines provided by domain-experts, we further add a post-processing step that explicitly enforces an increasing monotonicity constraint on the dose-response curve. Recall that by design, our metric learning approach learns to separate each pair of phenotypes with an equal distance. This proves to be problematic in a dose-response estimation context as in practice, our fungus can manifest more than two phenotypes along the dose gradient, thereby leading to oscillations. To circumvent this artefact, we leverage the Isotonic Regression technique using the Pool Adjacent Violators algorithm [7], which performs a least-squares fit while imposing a monotonicity constraint. We show an example curve that violates the latter constraint and its corrected version in Fig. 4.

**Figure 4.**
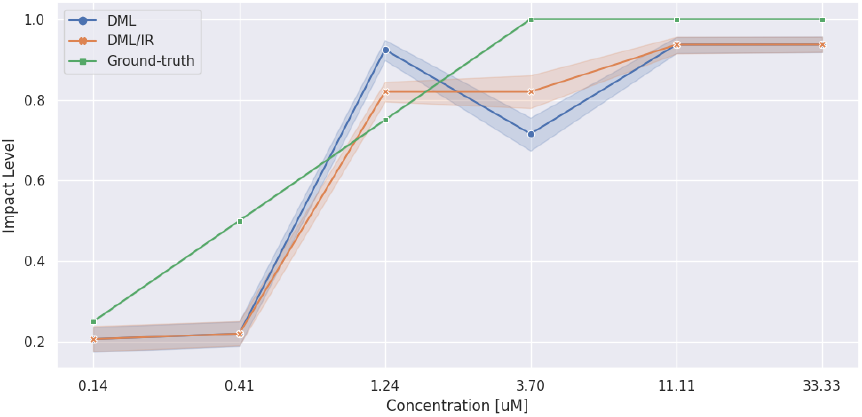
Example dose-response curve where isotonic regression is applied.

#### 3.2.2 Network architecture and Implementation

For the feature extraction function *f*_*θ*_, we leverage the popular ResNet50 architecture [11] for all image modalities. Next, each category-specific module 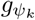 is of the form (BatchNorm, Dense(D, 256), ReLU, Dense(256, 256), ReLU, Dense(256, 128)), where BatchNorm is a batch normalization layer [12], ReLU is a rectifier linear unit activation function, and Dense(M,N) is a dense layer of input and output size M and N, respectively.

A specificity of cell painting image data is that intensity distributions show important discrepancies across different conditions, i.e. some conditions induce very strong fluorescence responses, while others, conversely, induce very weak responses. This scenario has shown to be problematic in the training phase, by bringing strong biases to samples of high intensities and thereby generating noisy gradients. We circumvent this issue by applying Adaptive Histogram Equalization [20] on each image to bring intensity distributions to the same range before feeding the image in the feature extraction backbone. Next, as the latter step discards biologically relevant information, we compute intensity statistics on each original image and use these further down in our pipeline. In particular, we concatenate global image statistics at the bottleneck level. As statistics, we use the 10-th and 90-th percentiles, the minimum and maximum intensity, and the standard deviation on all pixel values. Last, the scaling parameters *κ*_*l*_ are learned jointly using the output of *f*_*θ*_ through a dense layer.

#### 3.2.3 Training Procedure

We train the feature extraction backbone, the category-specific dense modules, the phenotypic-specific centroids, and the scaling parameters in an end-to-end fashion through stochastic gradient descent using the Adam optimizer [15]. At each iteration, we build a batch of 16 images from our training set, and perform mean-aggregation of crossentropy losses on each category modules. To account for imbalance in the number of images per phenotype, we sample a batch using weighted random sampling, i.e. scarcer phenotypes are sampled more often. As data augmentation, we select horizontal and vertical mirroring, as well as rotations in *{*90, 180, 270*}* degrees.

We train for 150 epochs with 200 iterations each. Due to memory constraint, we resize the input images from 2160 *×* 2160 to 512 *×* 512. The learning rate is initialized to 10*e*^*−*4^ and reduced to 10*e*^*−*5^ after 100 epochs. For validation, we retain 30% of images for each phenotype and compute on each category the balanced accuracy score using Eq. 3 after every epoch, and select the model that maximizes the average score on all categories.

## 4. Experiments

### 4.1. Baselines

#### 4.1.1 Fungi-Profiler: Hand-crafted Feature Extractor

Following the state-of-the-art in profiling of cell painting image data, we implement a feature extraction pipeline inspired by Cell-Profiler [25]. While the latter is designed for images of mammalian cells, where cells are well-defined and compact regions, we showed on Fig. 2 that filamentous fungi are quite different in nature, as the object of interest is generally non-compact following mycelium germination. This therefore prohibits the use of Cell-Profiler since it relies on a segmentation step that isolates individual cells prior to computing features.

We rather implement our own handcrafted feature extraction pipeline and name it *Fungi-Profiler*. As segmentation step, we perform a serie of edge-detection and morphological operations to isolate both spores and mycelium from the background. Next, we compute *∼* 210 features on each image. These are selected to encode, among others, general morphological properties, texture patterns at varying spatial scale, pixel intensity distributions, as well as cross-channel correlation information.

Finally, we implement a metric learning model similar to **DCML**, by replacing the deep feature extraction backbone with Fungi-Profiler features, and name it **FPCML**. The training procedure, calibration, and hyper-parameter selection are identical to **DCML**.

#### 4.1.2 Growth-Inhibition: Dose-response using Fungal Growth

Inspired by [24], we implement a simple baseline for doseresponse estimation that relies on fungal growth. In this approach, we assume that a biologically active compound inhibits the fungal growth, i.e. decreases the fungal mass as compared to a negative control condition, where spores germinate normally so as to form a mycelium.

Thus, we use our training set to compute *m*_0_ and *m*_1_, the average fungal masses that correspond to full growth (phenotype *FullGerm*) and no growth (phenotype *GermNone*), respectively, by using the same segmentation procedure used in *Fungi-Profiler*. We then optimize a linear interpolation model to map *m*_0_ to 0, and *m*_1_ to 1.

### 4.2. Test Dataset

We acquire cell painting image data on filamentous fungi *Botrytis cinerea* using 19 compound-treatments spanning 15 different biological targets, all pertaining to modes of actions described by the FRAC [9] through 2 plates (96 wells, 8 rows, 12 columns). Each compound is tested at 6 concentration values: 0.41 to 100*μM* in a dose-response range (dilution step 1*/*4), and images are acquired after a 24 hrs incubation time. To each plate, we further add a half row (6 wells) of negative-controls. We configure our microscope to capture 6 sites per well, i.e. non-overlapping regions.

### 4.3. Dose-response Curve Estimation

We now assess the performance of our metric learning methods in the context of dose-response curve estimation. Groundtruth impact-levels were assessed by a domain expert following visual inspection of image data. For practical reasons, we constrain values within the finite set [0, 0.25, 0.50, 0.75, 1], where 0 means no impact, and 1 means that the observed phenotype is similar to an annotated phenotype.

We use as performance metric the Mean Absolute Error (MAE) over the whole range of tested concentrations, and report that our best performing method, **DCML**, gives, averaged on all tested compounds, an MAE of 0.22, 0.24, 0.12, and 0.11 on apical cell wall, cell wall, lipid, and morphology, respectively. Next, we report substantial improvements w.r.t. **FPCML** on all categories, with a 19%, 35%, 14%, and 7% improvement on apical cell wall, cell wall, lipid, and morphology, respectively. Also, looking at the standard deviation of the MAE over all compounds, we note that method **DCML** provides much more robust estimates compared to baseline methods.

Concerning the growth-inhibition baseline, we note that it performs much worse than other methods overall, with an average MAE of 0.19, and in particular for conditions where the morphological impact is largely independent on the fungal mass, e.g. *GermAgglo*.

Finally, we show in Fig. 5, a subset of dose-response curves on all categories with associated groundtruths on all phenotypical categories.

**Figure 5.**
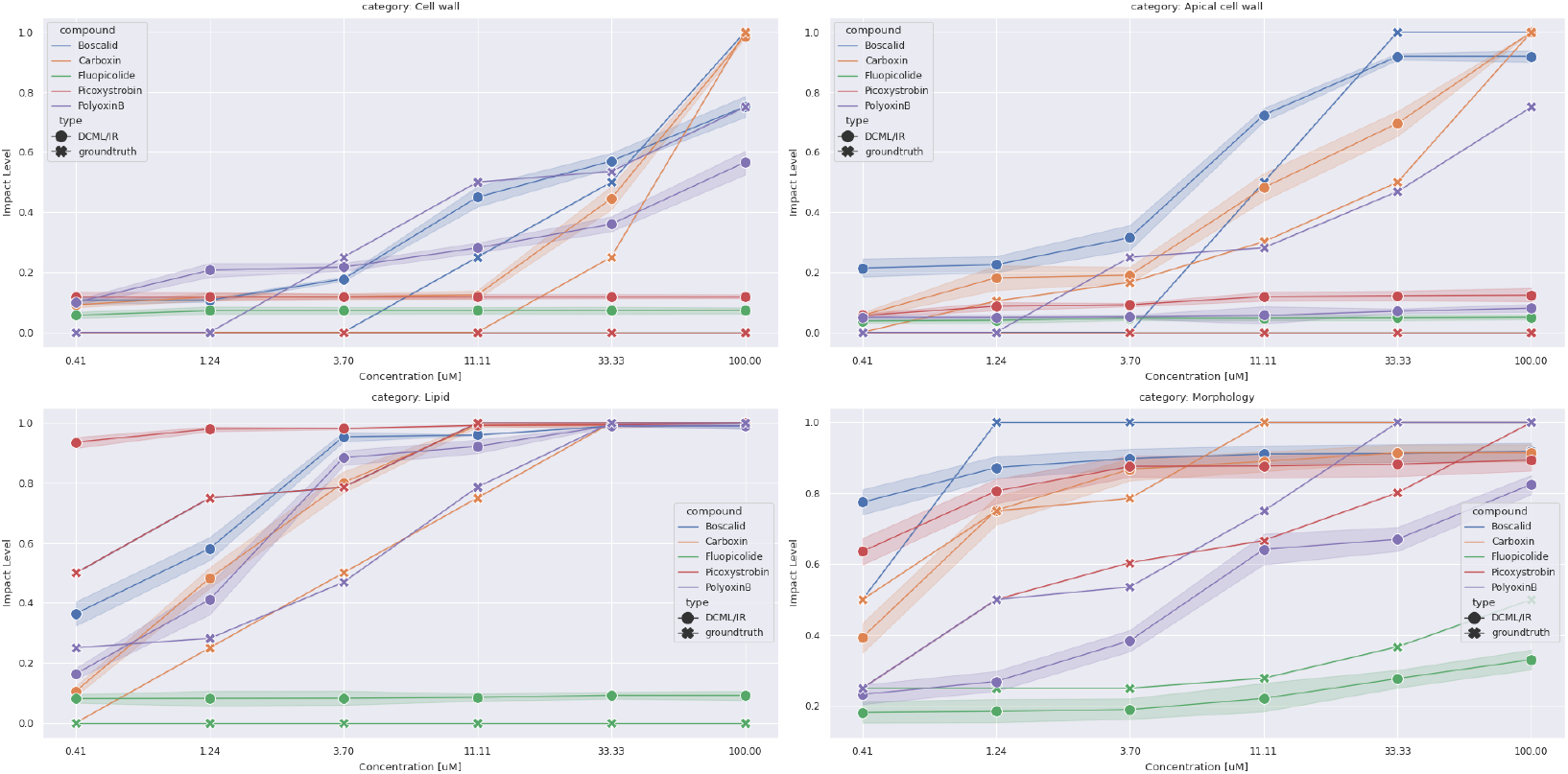
Example Dose-response curves on all phenotypical categories. We show estimates obtained with the proposed DCML method and the corresponding groundtruth values. Shaded regions correspond to the 95-th confidence interval on the mean.

### 4.4. Interpretable Cluster Analysis for Target Identification

Cluster analysis is a popular approach by which domain-experts look to test hypotheses regarding a new sample population by leveraging pairwise similarities with respect to another. In this section, we demonstrate the relevance of the proposed metric learning approach in this context. Recall that by design, our model learns metric-spaces specifically tailored to cluster phenotypes across different categories in an independent manner. Furthermore, given two populations of different compound and/or concentrations, similarities can be computed by combining distances pertaining to different phenotypical categories. Concretely, we leverage calibrated distances obtained with Eq. 5 and aggregate these using the mean operator to obtain a composite similarity metric.

We first select among all conditions of our test dataset (Sec. 4.2) those that are biologically active. Concretely, we leverage our dose-response estimations and pick conditions such that its response is above 0.75 according to at least one category. Next, we give in Fig. 6 visualization maps obtained through the t-SNE method [26] using pairwise distances obtained with Eq. 5. This analysis demonstrates the added value of our framework, and in particular how the phenotypes captured using the cell painting protocol are useful to separate many biological targets and modes of action, many of which are hardly separated using traditional microscopy. Last, we perform an ablation study where we quantify the clustering consistency on the same dataset using different combinations of categories. Concretely, we label each image-stack using the corresponding biological target or modes of action, and compute the mean silhouette scores over all image-stacks [22]. We report results using methods **DCML** and **FPCML** in Tab. 3.

**Figure 6.**
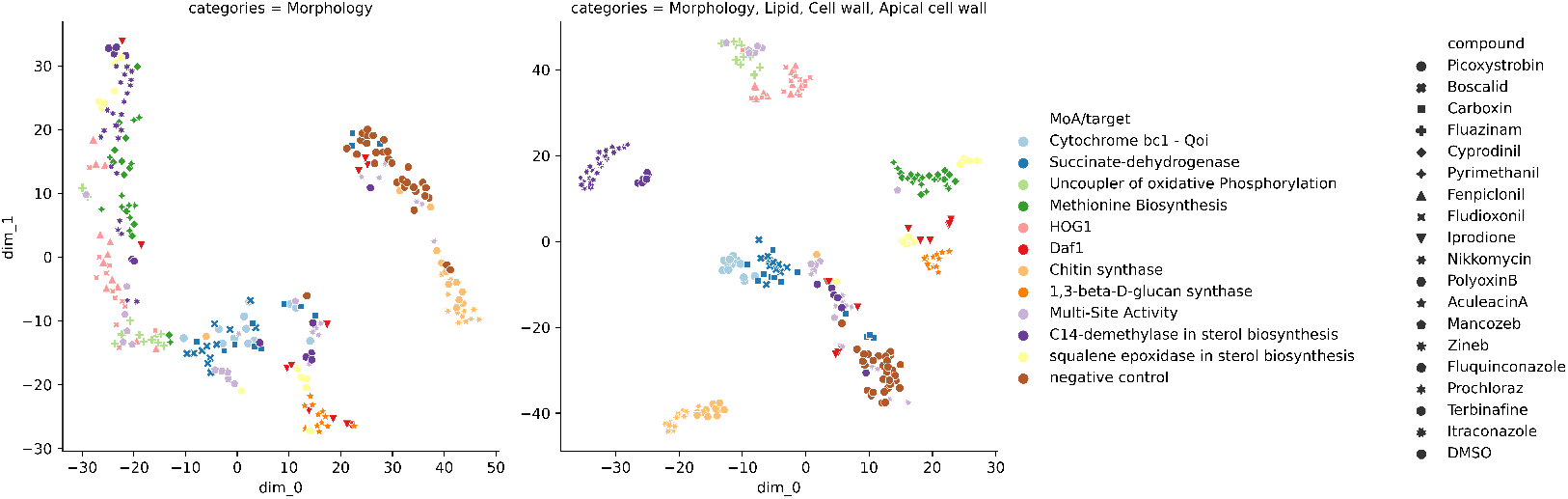
T-SNE visualization with different combinations of phenotypical categories. (Left) Pairwise distances using “Morphology” category. (Right) Pairwise distances obtained by mean-aggregation across all phenotypical categories. In the legend, colors indicate modes of action or biological target (when applicable), while symbols indicate compounds. Compound DMSO denotes negative controls.

**Table 3.** Silhouette scores of clustering w.r.t. biological targets for different combinations of phenotypical categories.

| Cell wall | Lipid | Apical cell wall | Morphology | DCML | FPCML |
| --- | --- | --- | --- | --- | --- |
| X | X | X | ✓ | -0.05 | -0.02 |
| X | X | ✓ | ✓ | 0.00 | 0.10 |
| X | ✓ | X | ✓ | 0.20 | 0.10 |
| X | ✓ | ✓ | ✓ | 0.23 | 0.06 |
| ✓ | X | X | ✓ | 0.07 | 0.05 |
| ✓ | X | ✓ | ✓ | 0.03 | 0.01 |
| ✓ | ✓ | X | ✓ | **0.25** | **0.12** |
| ✓ | ✓ | ✓ | ✓ | 0.22 | 0.09 |

Our best results are obtained using method **DCML**, further emphasizing the superiority of CNNs over hand-crafted features (**FPCML**). Also, we confirm the intuitions obtained in the previous experiments by noting a drastic increase in biological target clustering consistency when combining traditional microscopy with cell painting. In particular, we improve the silhouette score from *−*0.05 (morphology category only) to 0.25 (morphology, cell wall, and lipid categories). Interestingly, we note a decrease of consistency when combining both cell wall and apical cell wall categories, compared to using only one of them.

Finally, we note that among all available intra-cellular phenotypes, the lipid category is the most discriminative, followed by cell wall.

## 5. Conclusions

We contributed a framework that leverages cell painting image data, along with a Deep Cosine Metric Learning model to characterize the biological impact of compound treatments on filamentous fungi. Through a principled data exploration and annotation of phenotypes, we learned a set of metrics that allow to measure phenotypical transformations.

The proposed framework showed promising results in the task of estimating dose-response curves across a broad variety of phenotypes, both at a whole-cell and intra-cellular level. Furthermore, we showed that the learned metrics can be used to produce separable and combinable distances, thereby allowing domain experts to assess the importance of each phenotypical category in a cluster analysis setting, and gain precious insights into the biological impacts of compounds under test.

We further emphasize that even though machine learning models often exhibit performance levels on-par with humans, this frequently overshadows the problem of interpretability, where the model outputs arise from complex underlying operations that are hard, if not impossible, to understand, both for machine learning practitioner and domain experts. We believe that the proposed framework, to some extent, circumvents these limitations by providing distances that are easy to interpret.

As future perspectives, we aim to extend the current protocol to include additional fluorescent dyes, which, coupled with relevant phenotypical annotations, could potentially improve the understanding of fungi to compound treatments interactions.

## Supplementary Materials

### 1 Deep Cosine Metric Learning (DCML)

#### Algorithm 1: Training DML for one iteration

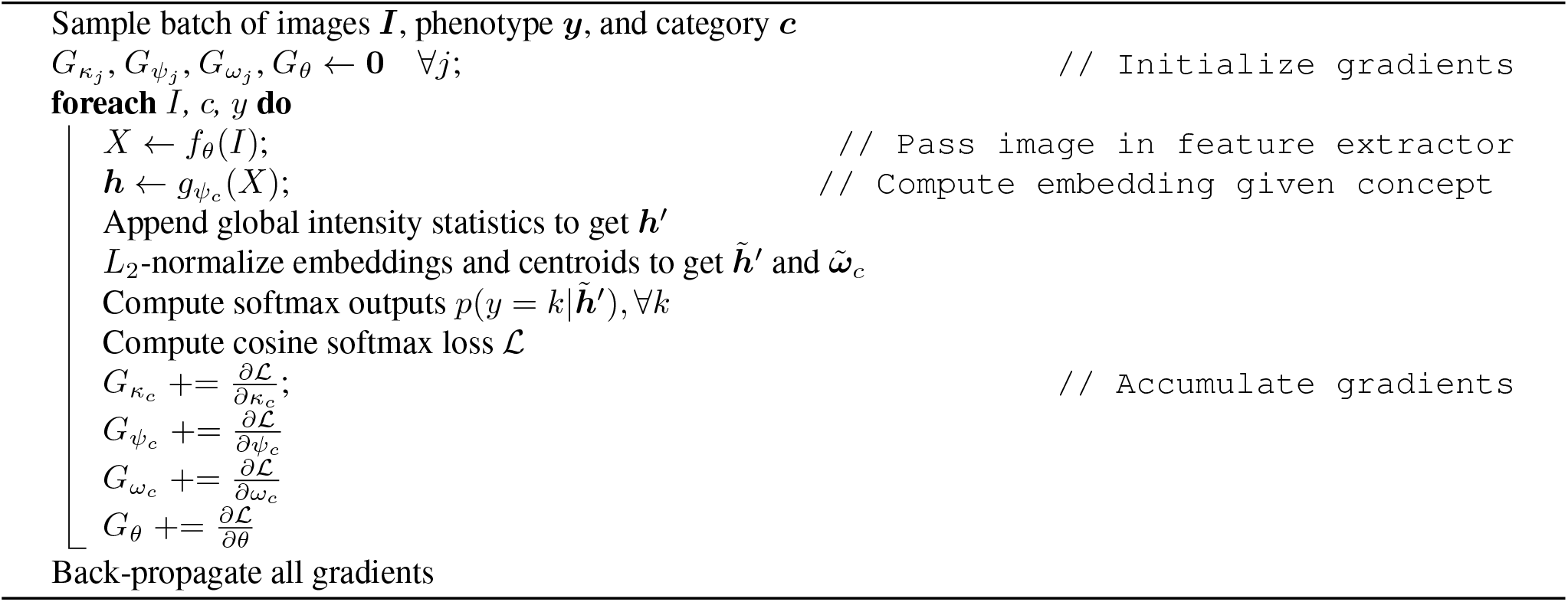

Sample batch of images ***I***, phenotype ***y***, and category ***c***

We show in Fig. 1 a schema of our architecture, and give a training pseudo-code in Alg. 1

### 2 Fungi-Profiler

We give here additional details related to *Fungi-Profiler*, our handcrafted feature extractor.

#### 2.1 Segmentation

As a first pre-processing step, We aim to segment regions where fungi is located to focus our downstream feature extraction pipeline.

We first apply Adaptive Histogram Equalization [7] on bright-field images to reduce luminosity variations accross different experiments. Next, we detect edges by applying an adaptive thresholding step, and follow with a serie of morphological operations. We show an example in Fig. 2.

**Figure 1:**
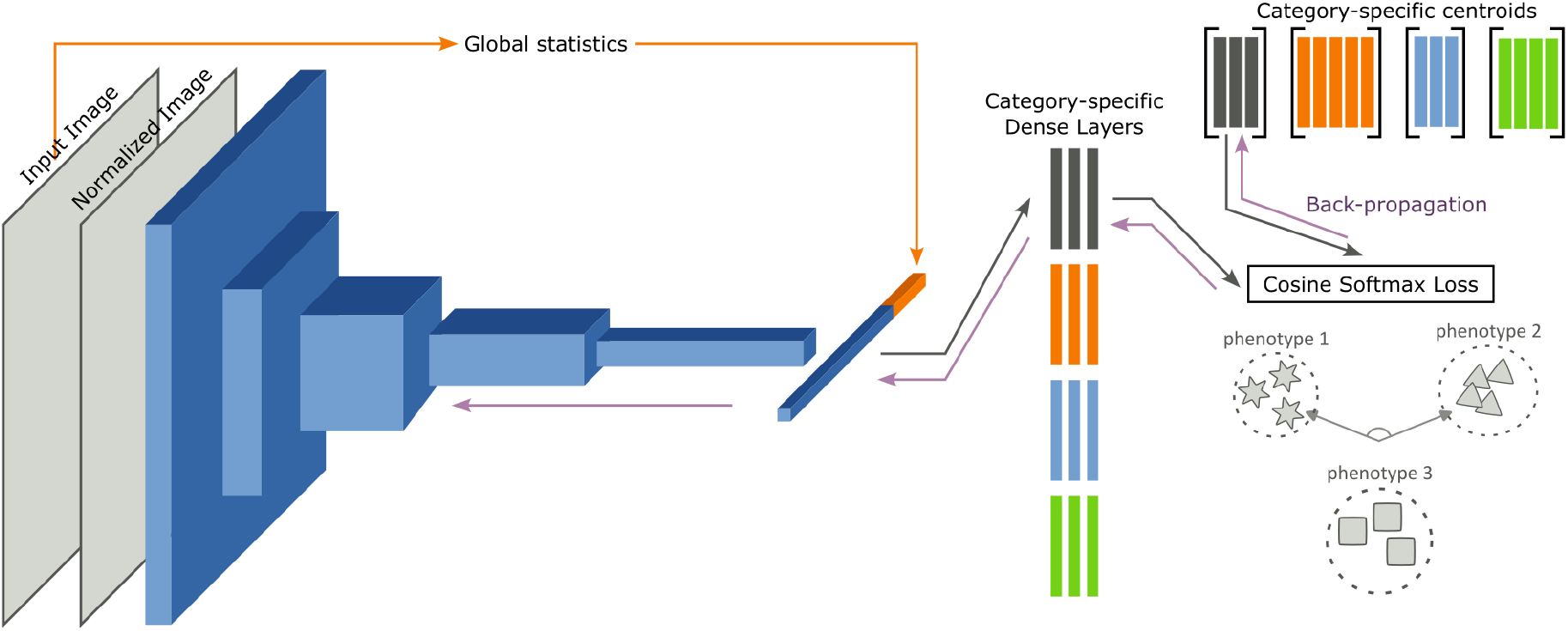
Deep Cosine Metric Learning model. Images are normalized, then fed into a Convolution Neural Network. We concatenate global image statistics computed on the original image to the resulting feature vector and feed it into a serie of dense layers according to the phenotypical category captured by its modality (red, green, blue and grey blocks). Last, we compute the cosine softmax loss on each image of a batch, and aggregate gradients to update the centroids, the category-specific dense layers, and the feature extraction backbone.

**Table 1:**
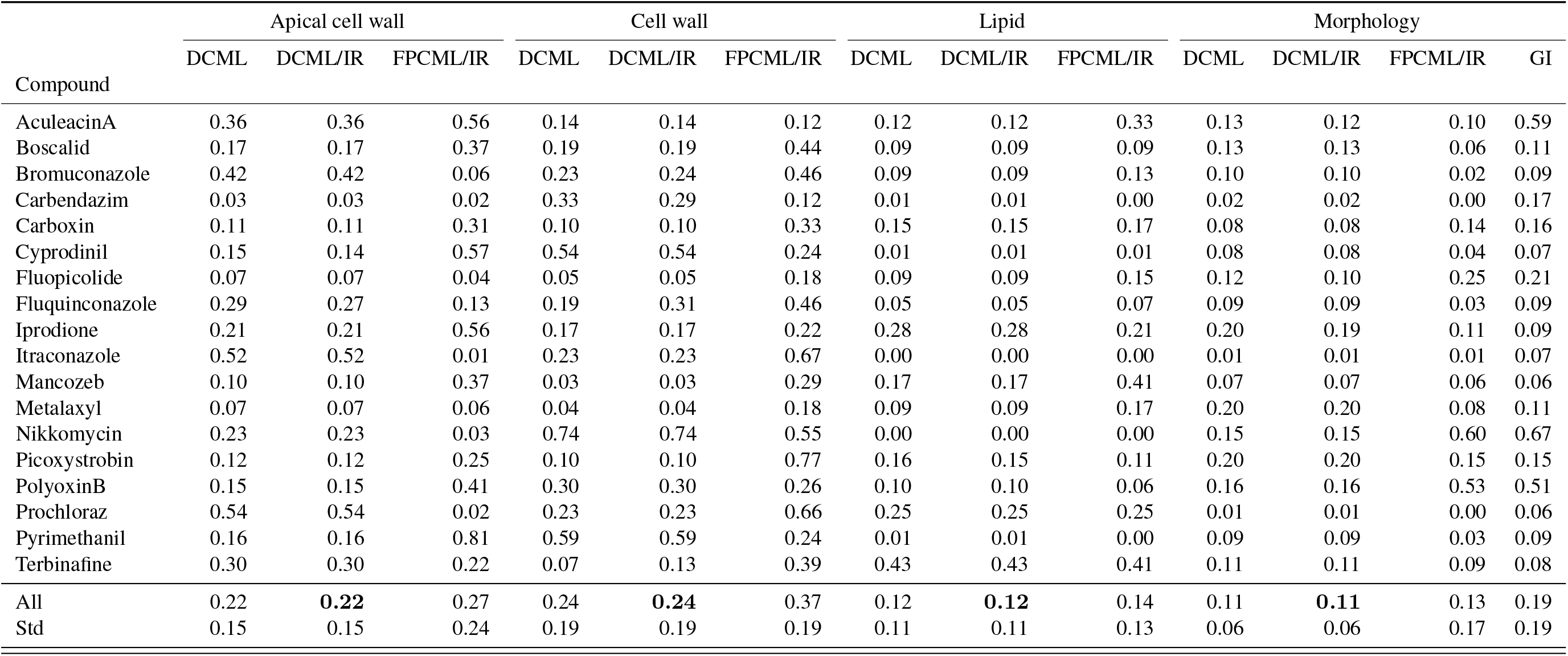
Mean absolute errors on each phenotypical categories, compound, and methods. DCML: Proposed deep cosine metric learning method, DCML/IR: DCML combined with isotonic regression post-processing. FPCML: Metric learning based on fungi-profiler features. GI: Growth-Inhibition. In bold, we show the best performing method.

**Figure 2:**
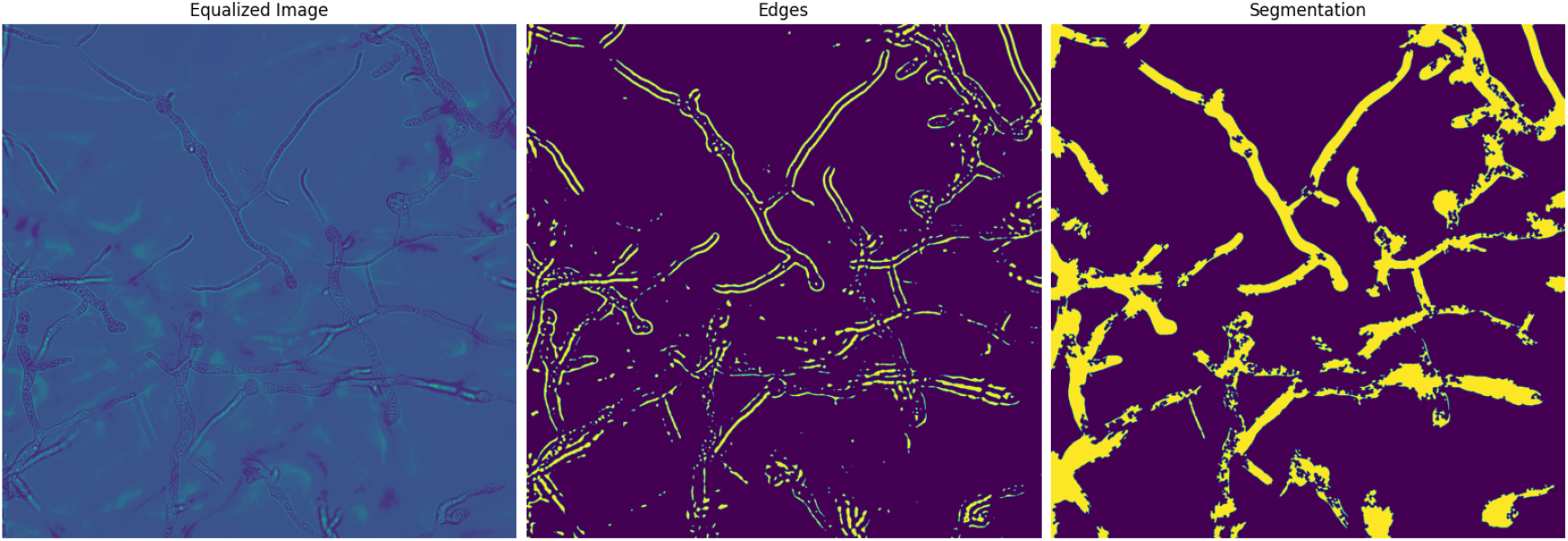
Segmentation step of Fungi-Profiler. (Left) Original image after adaptive histogram equalization. (Center) Edge map obtained through morphological operations. (Right) Segmentation map.

**Table 2:**
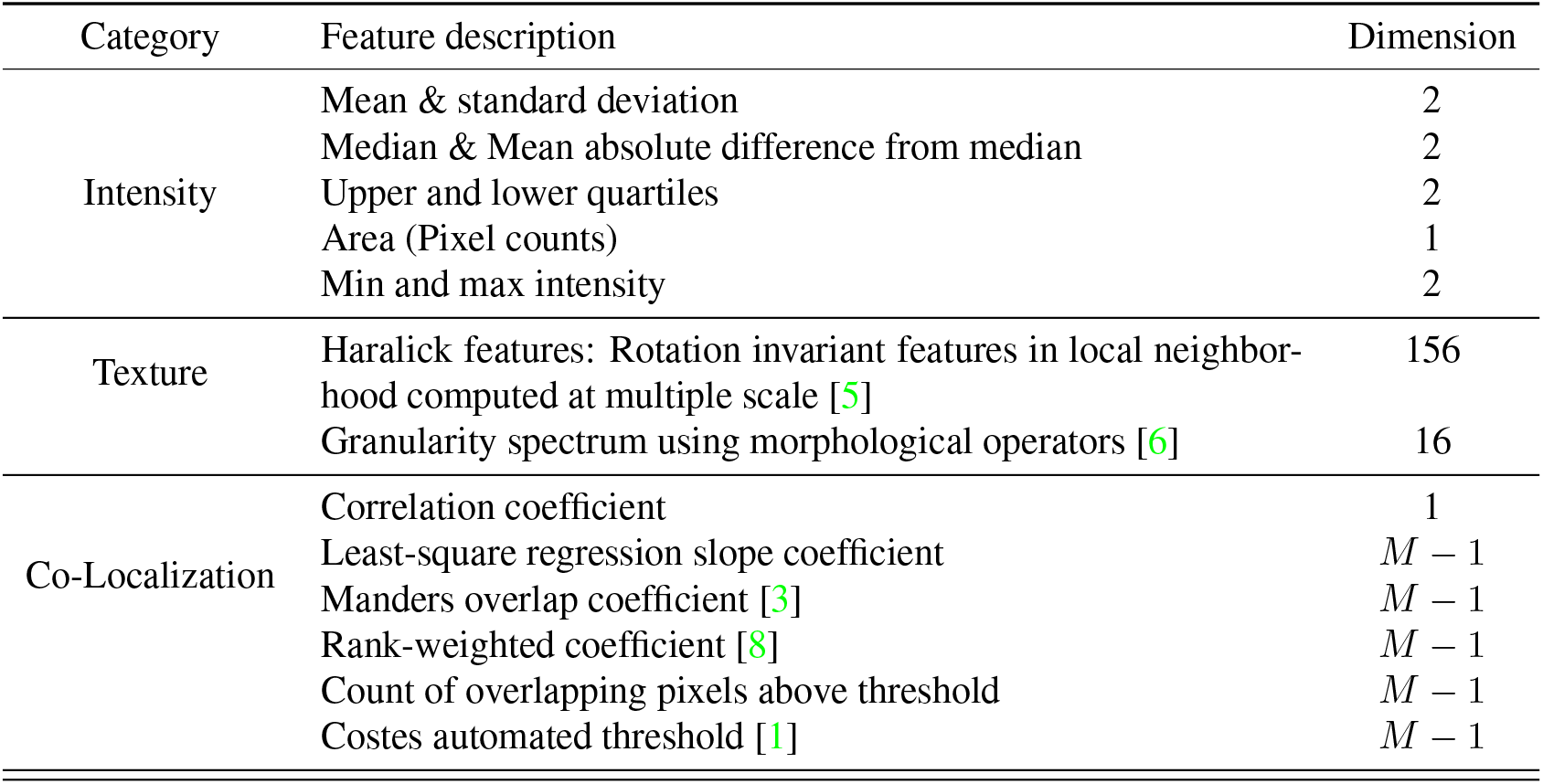
Summary of feature set computed by Fungi-Profiler. *M* denotes the number of image modalities.

| Category | Feature description | Dimension |
| --- | --- | --- |
| Intensity | Mean & standard deviation | 2 |
|  | Median & Mean absolute difference from median | 2 |
|  | Upper and lower quartiles | 2 |
|  | Area (Pixel counts) | 1 |
|  | Min and max intensity | 2 |
| Texture | Haralick features: Rotation invariant features in local neighborhood computed at multiple scale [5] | 156 |
|  | Granularity spectrum using morphological operators [6] | 16 |
| Co-Localization | Correlation coefficient | 1 |
| | Least-square regression slope coefficient | $M - 1$ |
| | Manders overlap coefficient [3] | $M - 1$ |
| | Rank-weighted coefficient [8] | $M - 1$ |
| | Count of overlapping pixels above threshold | $M - 1$ |
| | Costes automated threshold [1] | $M - 1$ |

#### 2.2 Illumination Correction

As noted in [9], microscopy images manifest an uneven illumination pattern accross the field of view, where regions close to the border show reduced brightness with respect to the center. This has been shown to have an important impact on the quality of visual features as measured by downstream prediction tasks.

To circumvent this artefact, we implement a simple illumination correction procedure as suggested by [9], where we compute an illumination correction function (ICF) as follows:

All bright-field images acquired during a plate assay are averaged pixel-wise, and then smoothed spatially using a squared median filter of size 25% of the original image size. Last, we apply the ICF to bright-field and fluorescent channels following:

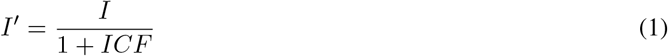

#### 2.3 Feature Extraction

In this step, we apply to each corrected image (both bright-field and fluorescent channels) the segmentation mask of the corresponding site, so as to set background pixels to 0. Next, we apply a serie of feature extraction routines summarized in Tab. 2 to obtain a vector of dimension 210 for each image. Note that, aside from traditional features computed on each image taken individually, we further leverage cross-channel information by extracting co-localization features on each pair of images of a given stack.

### 3 Pooled-Adjacent Violators Algorithm (PAVA)

We give in Alg. 2 a pseudo-code of the PAVA algorithm [2] by which we impose a monotonicity constraint on our dose-response estimations.

### 4 Testing Dataset

We show in Tab. 3 all compounds used in our experiments, along with their modes of action and biological targets.

#### Algorithm 2: Pooled-Adjacent Violators Algorithm (PAVA)

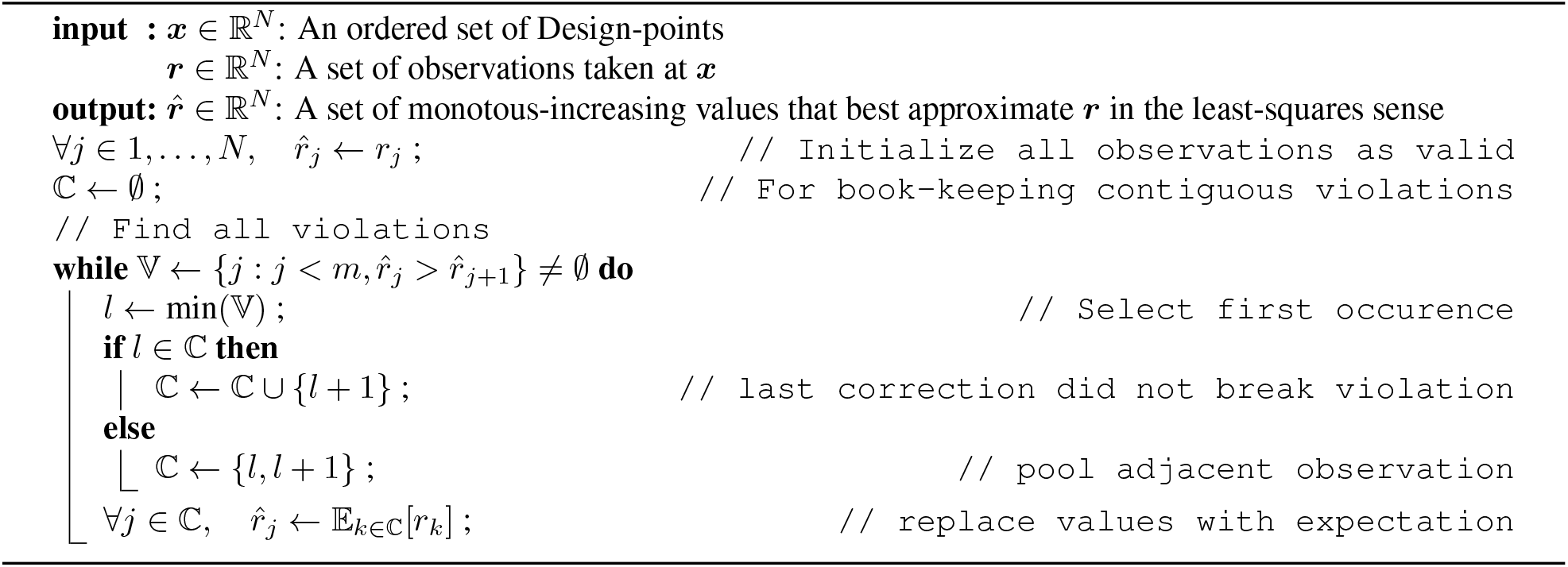

**Figure 3:**
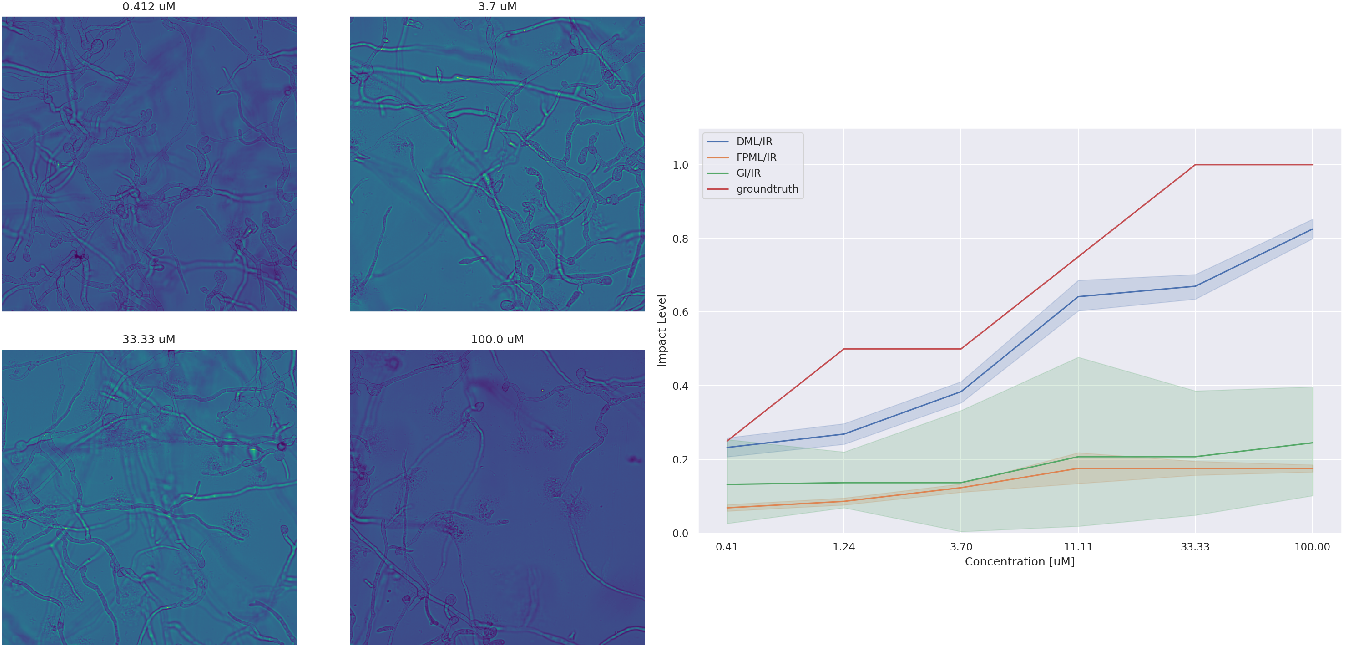
Example of a subtle phenotypical variation where spores turn bigger as concentration increases, while fungal mass is largely unaffected. Methods that relie on fungal mass (FPCML, GI) fail to capture this transformation, in contrast with DCML, which relies on a CNN feature extraction backbone.

### 5 Experiments

#### 5.1 Dose-response Estimation

We give in Tab. 1 the mean absolute errors of our methods on all tested compounds. In Fig. 3, we show a set of images along with estimated dose-response curves where a phenotype is largely independent on the fungal mass. In particular, we emphasize a limitation of methods **FPCML** and **GI** in capturing subtle variations in morphology, while our best method **DCML**, as it relies on a CNN backbone, behaves as expected.

---

**Table 3:**
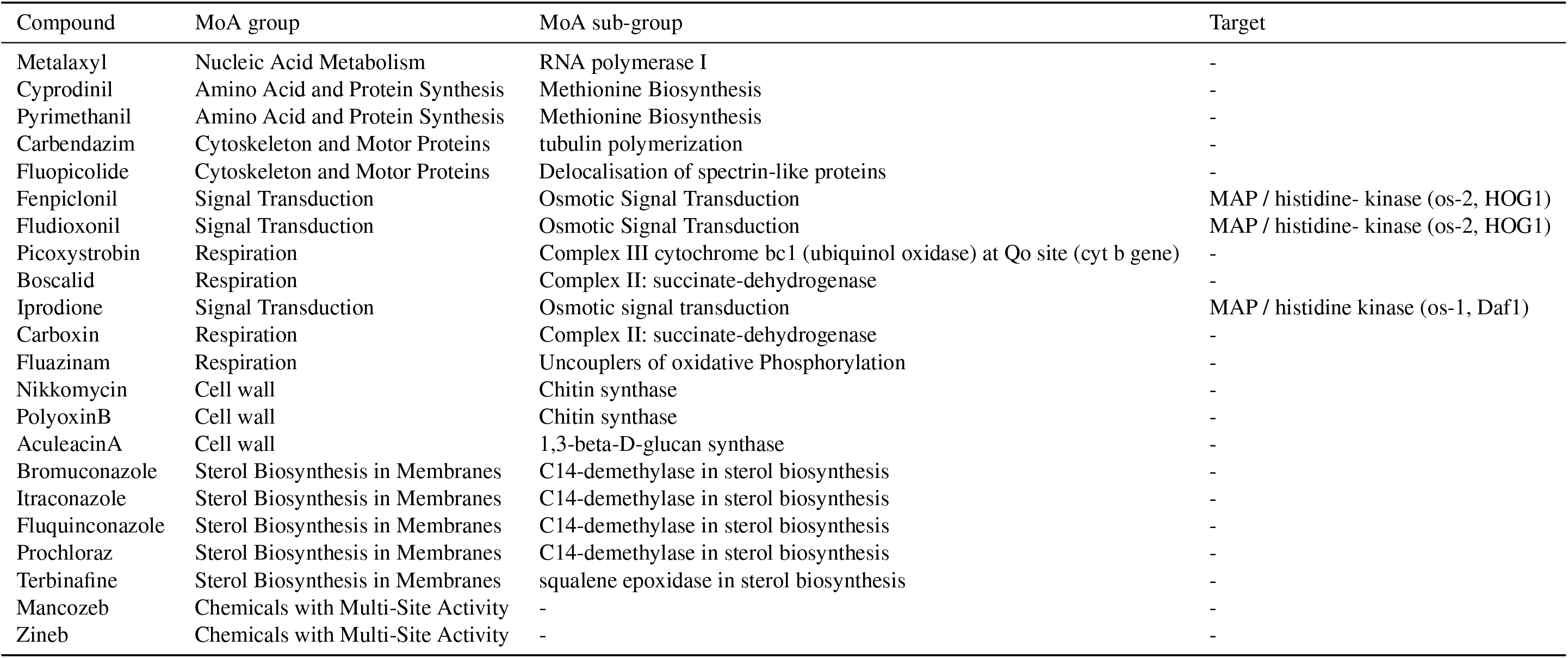
Summary of testing dataset. We show compound names, MoA group, MoA sub-group, and biological target according to the FRAC classification when applicable [4].

## Footnotes

1 For reproducibility, we make our dataset and code publicly available at https://doi.org/10.5281/zenodo.8227399.

## Notes

https://doi.org/10.5281/zenodo.8227399

### Competing Interest Statement

The authors have declared no competing interest.

